# An Interpretable Deep Learning Framework for Biomarker Discovery in Complex Disease Survival Outcomes

**DOI:** 10.1101/2025.09.30.679415

**Authors:** Shiyu Wan, Xinlei Mi, Fei Zou, Baiming Zou

## Abstract

Identification of important biomarkers associated with complex disease survival outcomes is fundamental for gaining an in-depth understanding of disease mechanisms and advancing precision medicine in conditions such as cancer and cardiovascular disorders. However, these tasks are complicated by the unique nature of time-to-event data, which captures both the occurrence and timing of clinical events. Notably, complex associations such as the non-linear and non-additive biomarker interactions and the high-dimensionality challenge conventional survival data modeling approaches. To address these difficulties, we propose SurvDNN, an enhanced deep neural network framework specifically designed for survival outcomes modeling. SurvDNN incorporates a bootstrapping-based regularization strategy to mitigate overfitting and a novel stability-driven filtering algorithm to improve model robustness. To enable interpretable biomarker discovery, we extend the Permutation-based Feature Importance Test (PermFIT) to survival settings, allowing rigorous quantification of individual biomarker contributions under complex biomarker–outcome associations. Through extensive simulations and applications to real-world datasets, SurvDNN consistently outperforms existing machine learning approaches in both biomarker identification and predictive accuracy. Our results demonstrate the potential of SurvDNN coupled with PermFIT as an interpretable, robust, and powerful tool for biomarker-driven survival modeling in complex diseases. An open-source R package implementing SurvDNN is publicly available on GitHub (https://github.com/BZou-lab/SurvDNN).

## 1 Introduction

Time-to-event data, commonly observed in clinical and medical practice, provide valuable insights into disease progression, treatment effectiveness, and patient outcomes over time [1, 2]. Accurate prediction of these outcomes yields crucial prognostic information, enabling healthcare professionals to identify individuals at heightened risk of adverse events. Such identification facilitates timely follow-up, targeted interventions, and personalized care optimized for efficacy while minimizing side effects [3]. For example, in oncology, precise survival outcome predictions guide decisions regarding surgical interventions, chemotherapy, and radiotherapy and play a key role in determining treatment intensity and duration [4]. Furthermore, time-to-event data enable continuous monitoring of disease progression, which is critical for adjusting treatment plans and managing patient expectations.

However, time-to-event data have unique characteristics that distinguish them from other data types and necessitate specialized modeling techniques. For instance, censoring occurs when the event of interest (e.g., death, failure, recovery) has not occurred by the end of the study or when a participant is lost to follow-up. Proper handling of censored observations is essential for valid inference and accurate survival risk prediction. Various analytical methods have been developed to address different types of time-to-event data and complex association structures. Parametric models, such as the accelerated failure time (AFT) models, have been essential in modeling survival data by assuming a particular underlying distribution for event times [5]. Semi-parametric methods, like the Cox proportional hazards model [6], are popular due to their flexibility in modeling hazard functions with an unspecified baseline hazard. However, accurately predicting time-to-event outcomes in clinical studies presents several additional analytic challenges. One significant challenge stems from the complex nonlinear and non-additive interactions among patient demographics and other biological and clinical risk factors, including molecular biomarkers, genetic variants, environmental exposures, and lifestyle factors. For instance, a multicenter observational study showed that the interactions among human epidermal growth factor receptor 2 expression levels, chemotherapy response level, and disease stage significantly affect the hazard in patients with gastric cancer [7]. Such intricate associations among these factors in affecting time-to-event outcomes can be challenging to model with parametric or semi-parametric models. This highlights the importance of using more flexible machine learning techniques in survival analysis [8, 9]. Integrating these methods with real-world clinical data, such as electronic health records and genomic data, can improve prediction accuracy and support clinical decision-making and patient care.

To mitigate the restrictive modeling assumptions (e.g., linearity, additivity) of parametric and semi-parametric survival models, a variety of machine learning methods have been introduced. Specifically, support vector machine (SVM) survival models utilize kernel functions, such as polynomial and radial basis function kernels, to capture intricate nonlinear relationships between predictors and survival outcomes [10, 11]. Moreover, tree-based machine learning methods, including random survival forest (RSF) [12] and extreme gradient boosting (XGBoost) [13], have been applied to survival data. These methods leverage decision trees as their building blocks for capturing complex non-linear relationships between predictors and time-to-event outcomes. In addition to these machine learning techniques, deep neural networks (DNNs) have also been employed for constructing predictive survival models, e.g., DeepSurv and DeepHit [14, 15]. The advantage of DNNs lies in their ability to robustly model complex functional relationships through multiple layers of nonlinear transformations. However, traditional DNN algorithms can be unstable in finite sample settings, due to their random parameter initialization [16–18]. As a result, there is a pressing need for more robust DNN methods that can effectively tackle the issue of instability.

While machine learning methods can alleviate restrictive assumptions about the relationship between predictors and time-to-event outcomes without the need for explicit functional relationship specifications, they are often perceived as black-box models in practice. Unlike parametric and semi-parametric models, the importance of each feature in these black-box models is often evaluated without rigorous statistical inference [19]. This lack of interpretability introduces another practical challenge for DNNs in survival analysis [20, 21]. Identifying prognostic features for survival outcomes is practically important, providing insight into disease mechanisms while improving risk prediction. Beyond prediction, rigorously evaluating each individual biomarker’s impact on survival outcomes in high-dimensional, non-linear settings is critical for precision oncology. Clinicians rely on validated biomarkers to stratify risk, guide therapy, and prioritize enrollment in targeted trials. Significant efforts have been made to evaluate the feature importance of time-to-event data in machine learning algorithms [22], including permutation importance for RSF, XGBoost’s boosting information, and filter methods such as joint mutual information maximization (JMIM) [23]. However, the RSF’s permutation importance [24, 25] may artificially favor features with numerous possible split points during variable selection [26, 27]. Also, RSF’s subsampling algorithm for statistical inference requires model refitting, which can be computationally intensive. XGBoost’s importance metric lacks a statistical significance test. The JMIM method typically requires transforming the survival outcome into an un-censored continuous outcome by calculating Cox model-based martingale residuals, which might be challenging to implement with complex DNN models [22, 28].

In many clinical settings, accurate disease-outcome prediction is critical, and so is the reliable identification of factors that underlie disease mechanisms. In precision oncology, this means not only stratifying patient risk but also pinpointing actionable biomarkers to guide management. In hepatocellular carcinoma (HCC), *α*-fetoprotein (AFP) levels inform therapeutic intensity and surveillance intervals, while liver-fibrosis staging determines surgical candidacy and transplant eligibility [29, 30]. Drawing on the Surveillance, Epidemiology, and End Results (SEER) program - a nationwide, population-based cancer registry maintained by the U.S. National Cancer Institute, the proposed research is motivated by a survival study of HCC patients [31]. This population-based dataset offers untapped insights into critical factors influencing post-diagnosis survival in real-world oncology practice. By identifying key features such as tumor stage and size, underlying liver function, biomarkers and demographic characteristics, we can clarify the factors that drive patient prognosis. Accurately predicting survival outcomes for HCC patients supports treatment selection, resource allocation, and prioritization of high-risk patients, thereby informing precision-medicine strategies and improving the quality of oncology care. Prior studies indicate that both age and the severity of liver fibrosis exert complex, non-linear, and interactive effects on HCC survival [32, 33], posing challenges in modeling and analyzing this type of complex survival data for conventional models. To address these challenges, we propose a refined DNN-based survival model, termed SurvDNN, which robustly models the complex associations between risk factors and survival outcomes. It incorporates two strategies to enhance the stability of conventional DNN methods: a bootstrapping procedure to prevent potential model overfitting, and a filtering algorithm to significantly enhance the stability of traditional DNN methods under finite sample settings. While SurvDNN excels at modeling complex relationships, it does not inherently provide per-feature importance or uncertainty estimates. To address this, we incorporate PermFIT, a permutation feature importance test originally developed for seemingly black-box machine learning models [19], into SurvDNN. This extension allows us to evaluate the importance of each individual feature in survival prediction. Furthermore, with the identified important feature from PermFIT, we can retrain the SurvDNN model to potentially boost the prediction performance. We refer to the retrained model as PermSurvDNN to distinguish it from the original SurvDNN trained with all available features.

We hypothesize that PermSurvDNN will (i) accurately recover clinically actionable biomarkers, and (ii) outperform existing methods in both discrimination and calibration for complex survival data, thereby informing biomarker-driven precision medicine. The remainder of this paper is structured as follows. Section 2.1 is dedicated to extensive simulation studies in which we compare SurvDNN against competing methods across diverse data-generating settings. In Section 2.2, we apply SurvDNN and other competing methods to two practical survival datasets that have motivated this study, further demonstrating its superiority over other methods. A brief discussion can be found in Section 3. In Section 4, we detail the proposed SurvDNN framework, along with the permutation-based testing procedure that enables rigorous testing of feature importance for time-to-event outcomes.

## 2 Results

### 2.1 Numerical Experiments

#### 2.1.1 Simulation Setting

To evaluate the performance of the proposed SurvDNN framework for survival data, we conducted numerical studies under two scenarios that captured a wide range of data structure complexity. For these scenarios, we generated the feature matrix ***X*** to encompass both continuous and discrete covariates, designated as ***X*** = (***x***_**1·**_, · · ·, ***x***_***n*·**_)*′*, with *p*-dimensional random vector 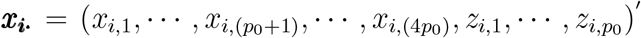, and *p*_0_ = 10, resulting in the total number of features *p* = 5*p*_0_ = 50. We denote the corresponding variables in ***X*** as *X*_*j*_ (*j* = 1, · ··, 4*p*_0_) and *Z*_*j*_ (*j* = 1, · · ·, *p*_0_), respectively, which correspond to the sets of continuous and discrete variables. Specifically, we let the continuous covariates 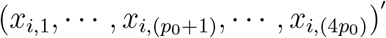 follow multivariate normal distribution *MV*ℕ (0, ∑), where ∑ = diag {∑_1_, ∑_2_, ∑_3_, ∑_4_}, a block-diagonal matrix with ∑_1_ = ∑_2_ = ∑_3_ = ∑_4_ being *p*_0_ × *p*_0_ compound symmetry matrices in which all their diagonal elements are equal to 1, and all off-diagonal elements are equal to *ρ*. Each of the discrete covariates 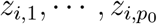 was independently drawn from the Bernoulli distribution with probability of 0.4. We considered various simulation settings with *ρ* ∈ {0, 0.3, 0.5}, *n* ∈ {1000, 3000}, and 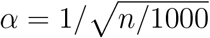 (a parameter used to maintain the covariate effects as sample size increased). Each simulation scenario was replicated 1,000 times.

##### Scenario 1

(Linear Effect): In this scenario, we considered a simple data-generating process in which all risk factors affect survival via linear effects only. Outcomes are generated from a proportional hazards model with a Gompertz baseline hazard, which is commonly used to model human mortality [34]:

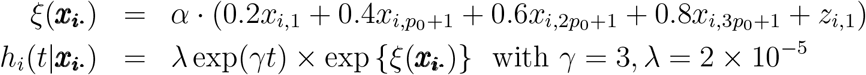

##### Scenario 2

(Non-linear Effect): In this scenario, we increased the data complexity from Scenario 1 by incorporating quadratic terms and interaction terms into *ξ*(·) as follows:

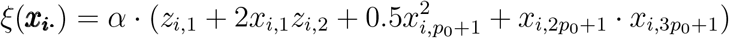

In all the scenarios, we independently drew the true event time *T*_true,*i*_ from the above settings using the R package *simsurv* [35], and the follow-up time *C*_*i*_ from a Weibull distribution (Shape = 1.5, Scale = 3.5). The observed outcome time for subject *i* was calculated as *T*_*i*_ = min(*T*_true,*i*_, *C*_*i*_), and the event indicator was generated as *δ*_*i*_ = 𝕀 (*T*_true,*i*_ ≤ *C*_*i*_) with 𝕀 being an indicator function. Finally, we combined these to obtain the simulated training dataset 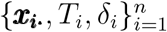. Testing datasets with sizes of 200 and 600 for training sizes of 1000 and 3000 were also generated to evaluate the performance of our proposed method and other statistical and machine-learning methods, corresponding to 80% vs 20% for training and testing samples setting.

### 2.1.2 Simulation Results

In the simulation studies, we compared SurvDNN with other competing methods, including Cox model, Lasso-Cox, RSF, XGBoost, DeepSurv and DeepHit. The implementation details of these methods are provided in the Appendix. The frequencies of the important features detected by each method are reported in Tables 1 and 2. Because PermFIT is a model-agnostic feature-importance method, we denote its application to a specific model as PermModel (e.g., PermSurvDNN, PermDeepSurv). For the PermFIT-adopted methods, original Cox model, and original RSF that provide *p*-values, we set the significance level to *α* = 0.05. For Lasso-Cox, features with non-zero regression coefficients were defined to be important. We define *S*_*I*_ as the set of truly important features. We employed 3 metrics to evaluate the feature selection performance, including average power, type I error, and average rank sum:

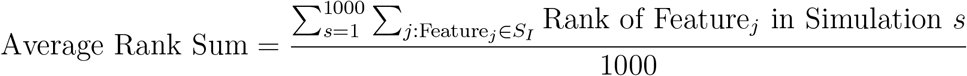

**Table 1:** Feature Importance Identification Performance Comparison under Scenario 1.

| Scenario 1a: $n = 1,000$ | | | | | | | | | | | | |
| --- | --- | --- | --- | --- | --- | --- | --- | --- | --- | --- | --- | --- |
| $\rho$ | Variable | Perm-SurvDNN | Perm-Cox | Perm-DeepHit | Perm-DeepSurv | Perm-RSF | Perm-XGBoost | AFT | Cox | Lasso-Cox | RSF | XGBoost |
| 0 | $X_1$ | 0.652 | 0.693 | 0.216 | 0.258 | 0.468 | 0.318 | 0.944 | 0.941 | 0.985 | 0.996 | / |
| | $X_{p_0+1}$ | 0.995 | 0.995 | 0.778 | 0.816 | 0.982 | 0.911 | 0.966 | 0.995 | 1.000 | 0.996 | / |
| | $X_{2p_0+1}$ | 0.995 | 0.995 | 0.995 | 0.998 | 1.000 | 1.000 | 0.964 | 1.000 | 1.000 | 1.000 | / |
| | $X_{3p_0+1}$ | 0.999 | 1.000 | 0.997 | 1.000 | 1.000 | 1.000 | 0.985 | 0.999 | 1.000 | 1.000 | / |
| | $Z_1$ | 1.000 | 1.000 | 0.828 | 0.881 | 0.996 | 0.974 | 0.970 | 1.000 | 0.997 | 0.998 | / |
|  | Avg Power | 0.928 | 0.936 | 0.763 | 0.790 | 0.889 | 0.841 | 0.966 | 0.987 | 0.996 | 0.999 | / |
|  | Type I Error | 0.044 | 0.044 | 0.054 | 0.051 | 0.053 | 0.053 | 0.924 | 0.071 | 0.240 | 0.887 | / |
|  | Avg Rank Sum | 18.39 | 17.24 | 31.36 | 30.16 | 21.34 | 26.31 | 89.15 | 15.71 | 15.64 | 21.67 | 21.11 |
| 0.3 | $X_1$ | 0.480 | 0.675 | 0.207 | 0.182 | 0.346 | 0.231 | 0.934 | 0.856 | 0.973 | 1.000 | / |
| | $X_{p_0+1}$ | 0.978 | 0.997 | 0.740 | 0.791 | 0.922 | 0.847 | 0.956 | 1.000 | 0.996 | 1.000 | / |
| | $X_{2p_0+1}$ | 1.000 | 0.999 | 0.985 | 0.982 | 1.000 | 0.994 | 0.964 | 1.000 | 0.997 | 1.000 | / |
| | $X_{3p_0+1}$ | 1.000 | 1.000 | 0.993 | 1.000 | 1.000 | 1.000 | 0.967 | 1.000 | 1.000 | 1.000 | / |
| | $Z_1$ | 0.996 | 0.997 | 0.884 | 0.870 | 0.998 | 0.984 | 0.970 | 0.996 | 1.000 | 0.999 | / |
|  | Avg Power | 0.891 | 0.934 | 0.762 | 0.765 | 0.853 | 0.811 | 0.958 | 0.970 | 0.993 | 1.000 | / |
|  | Type I Error | 0.038 | 0.069 | 0.069 | 0.061 | 0.042 | 0.063 | 0.925 | 0.074 | 0.238 | 0.890 | / |
|  | Avg Rank Sum | 20.34 | 18.46 | 32.62 | 33.74 | 23.57 | 29.30 | 98.98 | 16.80 | 16.08 | 27.12 | 23.60 |
| 0.5 | $X_1$ | 0.323 | 0.703 | 0.162 | 0.215 | 0.196 | 0.186 | 0.959 | 0.736 | 0.950 | 1.000 | / |
| | $X_{p_0+1}$ | 0.866 | 0.994 | 0.570 | 0.644 | 0.785 | 0.776 | 0.946 | 1.000 | 0.996 | 1.000 | / |
| | $X_{2p_0+1}$ | 1.000 | 0.999 | 0.930 | 0.965 | 0.995 | 0.986 | 0.949 | 1.000 | 0.997 | 1.000 | / |
| | $X_{3p_0+1}$ | 1.000 | 1.000 | 0.997 | 1.000 | 0.995 | 1.000 | 0.962 | 1.000 | 1.000 | 1.000 | / |
| | $Z_1$ | 0.996 | 0.997 | 0.785 | 0.910 | 0.996 | 0.984 | 0.970 | 0.996 | 1.000 | 1.000 | / |
|  | Avg Power | 0.837 | 0.939 | 0.689 | 0.747 | 0.793 | 0.786 | 0.957 | 0.946 | 0.989 | 1.000 | / |
|  | Type I Error | 0.035 | 0.094 | 0.079 | 0.087 | 0.038 | 0.070 | 0.928 | 0.070 | 0.238 | 0.895 | / |
|  | Avg Rank Sum | 23.84 | 19.65 | 42.81 | 36.42 | 27.77 | 32.88 | 99.63 | 18.37 | 16.65 | 35.25 | 26.72 |
| Scenario 1b: $n = 3,000$ | | | | | | | | | | | | |
| $\rho$ | Variable | Perm-SurvDNN | Perm-Cox | Perm-DeepHit | Perm-DeepSurv | Perm-RSF | Perm-XGBoost | AFT | Cox | Lasso-Cox | RSF | XGBoost |
| 0 | $X_1$ | 0.775 | 0.842 | 0.386 | 0.365 | 0.619 | 0.358 | 0.911 | 0.984 | 0.988 | 0.993 | / |
| | $X_{p_0+1}$ | 0.995 | 0.995 | 0.927 | 0.913 | 0.997 | 0.924 | 0.904 | 0.995 | 1.000 | 0.996 | / |
| | $X_{2p_0+1}$ | 0.995 | 0.995 | 1.000 | 0.998 | 1.000 | 1.000 | 0.913 | 1.000 | 1.000 | 1.000 | / |
| | $X_{3p_0+1}$ | 0.999 | 1.000 | 1.000 | 0.995 | 1.000 | 1.000 | 0.920 | 1.000 | 1.000 | 1.000 | / |
| | $Z_1$ | 1.000 | 1.000 | 0.997 | 0.988 | 0.996 | 0.993 | 0.919 | 1.000 | 0.997 | 0.998 | / |
|  | Avg Power | 0.953 | 0.966 | 0.812 | 0.852 | 0.922 | 0.855 | 0.913 | 0.996 | 0.997 | 0.997 | / |
|  | Type I Error | 0.046 | 0.042 | 0.056 | 0.050 | 0.058 | 0.055 | 0.905 | 0.056 | 0.227 | 0.744 | / |
|  | Avg Rank Sum | 16.99 | 16.28 | 25.16 | 24.39 | 19.40 | 25.22 | 101.28 | 15.40 | 15.37 | 20.62 | 24.69 |
| 0.3 | $X_1$ | 0.648 | 0.792 | 0.299 | 0.335 | 0.434 | 0.366 | 0.904 | 0.909 | 0.988 | 0.995 | / |
| | $X_{p_0+1}$ | 0.996 | 0.999 | 0.881 | 0.891 | 0.978 | 0.914 | 0.911 | 1.000 | 0.996 | 1.000 | / |
| | $X_{2p_0+1}$ | 1.000 | 0.999 | 0.996 | 0.994 | 1.000 | 0.993 | 0.906 | 1.000 | 0.997 | 1.000 | / |
| | $X_{3p_0+1}$ | 1.000 | 1.000 | 0.998 | 1.000 | 1.000 | 1.000 | 0.910 | 1.000 | 1.000 | 1.000 | / |
| | $Z_1$ | 0.996 | 0.997 | 0.981 | 0.993 | 0.998 | 1.000 | 0.912 | 0.996 | 1.000 | 0.999 | / |
|  | Avg Power | 0.928 | 0.957 | 0.831 | 0.843 | 0.882 | 0.855 | 0.909 | 0.981 | 0.996 | 0.998 | / |
|  | Type I Error | 0.041 | 0.054 | 0.068 | 0.061 | 0.038 | 0.069 | 0.902 | 0.054 | 0.226 | 0.763 | / |
|  | Avg Rank Sum | 18.31 | 17.03 | 28.83 | 26.52 | 21.16 | 26.14 | 105.79 | 15.88 | 15.66 | 24.54 | 25.66 |
| 0.5 | $X_1$ | 0.431 | 0.758 | 0.233 | 0.238 | 0.224 | 0.283 | 0.921 | 0.789 | 0.955 | 0.985 | / |
| | $X_{p_0+1}$ | 0.973 | 0.999 | 0.860 | 0.844 | 0.885 | 0.871 | 0.904 | 1.000 | 0.996 | 1.000 | / |
| | $X_{2p_0+1}$ | 1.000 | 0.999 | 1.000 | 0.960 | 1.000 | 0.993 | 0.899 | 1.000 | 0.997 | 1.000 | / |
| | $X_{3p_0+1}$ | 1.000 | 1.000 | 1.000 | 0.967 | 1.000 | 1.000 | 0.910 | 1.000 | 1.000 | 1.000 | / |
| | $Z_1$ | 0.996 | 0.997 | 0.969 | 0.957 | 0.998 | 0.991 | 0.919 | 0.996 | 1.000 | 0.999 | / |
|  | Avg Power | 0.880 | 0.951 | 0.812 | 0.793 | 0.821 | 0.827 | 0.911 | 0.957 | 0.990 | 0.997 | / |
|  | Type I Error | 0.038 | 0.092 | 0.089 | 0.077 | 0.038 | 0.094 | 0.900 | 0.057 | 0.237 | 0.786 | / |
|  | Avg Rank Sum | 21.28 | 18.79 | 31.41 | 28.57 | 25.32 | 31.37 | 109.37 | 17.66 | 16.50 | 32.31 | 30.53 |

**Table 2:** Feature Importance Identification Performance Comparison under Scenario 2.

| Scenario 2a: $n = 1,000$ | | | | | | | | | | | | |
| --- | --- | --- | --- | --- | --- | --- | --- | --- | --- | --- | --- | --- |
| $\rho$ | Variable | Perm-SurvDNN | Perm-Cox | Perm-DeepHit | Perm-DeepSurv | Perm-RSF | Perm-XGBoost | AFT | Cox | Lasso-Cox | RSF | XGBoost |
| 0 | $X_1$ | 1.000 | 1.000 | 0.945 | 0.983 | 1.000 | 1.000 | 0.993 | 1.000 | 1.000 | 1.000 | / |
| | $X_{p_0+1}$ | 0.947 | 0.075 | 0.167 | 0.165 | 1.000 | 0.995 | 0.599 | 0.130 | 0.119 | 1.000 | / |
| | $X_{2p_0+1}$ | 0.988 | 0.060 | 0.240 | 0.224 | 0.727 | 0.725 | 0.674 | 0.137 | 0.186 | 1.000 | / |
| | $X_{3p_0+1}$ | 0.980 | 0.054 | 0.241 | 0.198 | 0.698 | 0.712 | 0.657 | 0.116 | 0.174 | 1.000 | / |
| | $Z_1$ | 0.997 | 0.995 | 0.468 | 0.461 | 0.991 | 0.921 | 0.899 | 1.000 | 1.000 | 1.000 | / |
| | $Z_2$ | 0.993 | 0.252 | 0.194 | 0.121 | 1.000 | 1.000 | 0.764 | 0.274 | 0.205 | 1.000 | / |
|  | Avg Power | 0.984 | 0.406 | 0.376 | 0.359 | 0.903 | 0.892 | 0.764 | 0.443 | 0.447 | 1.000 | / |
|  | Type I Error | 0.049 | 0.043 | 0.051 | 0.055 | 0.048 | 0.056 | 0.658 | 0.080 | 0.121 | 0.874 | / |
|  | Avg Rank Sum | 21.84 | 103.87 | 79.61 | 88.44 | 26.43 | 28.41 | 121.24 | 96.02 | 104.49 | 27.07 | 30.88 |
| 0.3 | $X_1$ | 1.000 | 1.000 | 0.985 | 0.976 | 1.000 | 1.000 | 0.962 | 1.000 | 1.000 | 1.000 | / |
| | $X_{p_0+1}$ | 0.985 | 0.085 | 0.316 | 0.293 | 1.000 | 0.997 | 0.598 | 0.107 | 0.113 | 1.000 | / |
| | $X_{2p_0+1}$ | 0.997 | 0.074 | 0.345 | 0.312 | 0.750 | 0.735 | 0.673 | 0.103 | 0.166 | 1.000 | / |
| | $X_{3p_0+1}$ | 0.994 | 0.091 | 0.341 | 0.328 | 0.741 | 0.748 | 0.677 | 0.124 | 0.189 | 1.000 | / |
| | $Z_1$ | 0.998 | 0.995 | 0.590 | 0.535 | 0.994 | 0.941 | 0.885 | 0.997 | 1.000 | 1.000 | / |
| | $Z_2$ | 0.997 | 0.246 | 0.292 | 0.207 | 1.000 | 1.000 | 0.737 | 0.245 | 0.198 | 1.000 | / |
|  | Avg Power | 0.995 | 0.415 | 0.478 | 0.442 | 0.914 | 0.903 | 0.755 | 0.429 | 0.444 | 1.000 | / |
|  | Type I Error | 0.061 | 0.066 | 0.068 | 0.063 | 0.047 | 0.065 | 0.660 | 0.081 | 0.118 | 0.888 | / |
|  | Avg Rank Sum | 21.21 | 104.23 | 71.17 | 75.88 | 25.38 | 27.98 | 122.49 | 98.35 | 104.77 | 30.74 | 29.96 |
| 0.5 | $X_1$ | 1.000 | 1.000 | 0.977 | 0.955 | 1.000 | 1.000 | 0.964 | 1.000 | 1.000 | 1.000 | / |
| | $X_{p_0+1}$ | 0.983 | 0.098 | 0.547 | 0.485 | 1.000 | 0.998 | 0.651 | 0.075 | 0.080 | 1.000 | / |
| | $X_{2p_0+1}$ | 0.998 | 0.090 | 0.500 | 0.505 | 0.748 | 0.787 | 0.661 | 0.085 | 0.148 | 1.000 | / |
| | $X_{3p_0+1}$ | 0.998 | 0.100 | 0.486 | 0.475 | 0.743 | 0.812 | 0.679 | 0.095 | 0.160 | 1.000 | / |
| | $Z_1$ | 0.998 | 0.100 | 0.658 | 0.700 | 0.998 | 0.945 | 0.919 | 0.995 | 1.000 | 1.000 | / |
| | $Z_2$ | 1.000 | 0.292 | 0.481 | 0.405 | 1.000 | 1.000 | 0.781 | 0.248 | 0.238 | 1.000 | / |
|  | Avg Power | 0.995 | 0.429 | 0.608 | 0.588 | 0.913 | 0.924 | 0.776 | 0.416 | 0.438 | 1.000 | / |
|  | Type I Error | 0.090 | 0.100 | 0.099 | 0.100 | 0.055 | 0.086 | 0.673 | 0.076 | 0.127 | 0.898 | / |
|  | Avg Rank Sum | 21.37 | 104.42 | 57.72 | 60.60 | 25.80 | 26.52 | 122.00 | 100.22 | 105.75 | 39.20 | 30.41 |
| Scenario 2b: $n = 3,000$ | | | | | | | | | | | | |
| $\rho$ | Variable | Perm-SurvDNN | Perm-Cox | Perm-DeepHit | Perm-DeepSurv | Perm-RSF | Perm-XGBoost | AFT | Cox | Lasso-Cox | RSF | XGBoost |
| 0 | $X_1$ | 1.000 | 1.000 | 1.000 | 0.991 | 1.000 | 1.000 | 0.992 | 1.000 | 1.000 | 1.000 | / |
| | $X_{p_0+1}$ | 1.000 | 0.064 | 0.266 | 0.273 | 1.000 | 1.000 | 0.911 | 0.109 | 0.103 | 1.000 | / |
| | $X_{2p_0+1}$ | 1.000 | 0.049 | 0.467 | 0.438 | 0.990 | 0.987 | 0.946 | 0.087 | 0.177 | 1.000 | / |
| | $X_{3p_0+1}$ | 1.000 | 0.058 | 0.486 | 0.442 | 0.989 | 0.984 | 0.923 | 0.103 | 0.187 | 1.000 | / |
| | $Z_1$ | 1.000 | 1.000 | 0.795 | 0.800 | 1.000 | 0.983 | 0.966 | 1.000 | 1.000 | 1.000 | / |
| | $Z_2$ | 1.000 | 0.144 | 0.316 | 0.322 | 1.000 | 1.000 | 0.929 | 0.136 | 0.119 | 1.000 | / |
|  | Avg Power | 1.000 | 0.386 | 0.555 | 0.544 | 0.996 | 0.992 | 0.944 | 0.405 | 0.432 | 1.000 | / |
|  | Type I Error | 0.065 | 0.045 | 0.055 | 0.051 | 0.055 | 0.059 | 0.934 | 0.062 | 0.124 | 0.754 | / |
|  | Avg Rank Sum | 21.00 | 107.71 | 54.93 | 57.67 | 21.19 | 21.48 | 135.08 | 102.32 | 107.89 | 23.57 | 36.43 |
| 0.3 | $X_1$ | 1.000 | 1.000 | 0.995 | 0.993 | 1.000 | 1.000 | 0.987 | 1.000 | 1.000 | 1.000 | / |
| | $X_{p_0+1}$ | 1.000 | 0.079 | 0.503 | 0.554 | 1.000 | 1.000 | 0.931 | 0.072 | 0.083 | 1.000 | / |
| | $X_{2p_0+1}$ | 1.000 | 0.069 | 0.604 | 0.680 | 0.988 | 0.989 | 0.936 | 0.086 | 0.162 | 1.000 | / |
| | $X_{3p_0+1}$ | 1.000 | 0.070 | 0.574 | 0.711 | 0.992 | 0.988 | 0.921 | 0.078 | 0.170 | 1.000 | / |
| | $Z_1$ | 1.000 | 1.000 | 0.848 | 0.876 | 1.000 | 0.987 | 0.960 | 1.000 | 1.000 | 1.000 | / |
| | $Z_2$ | 1.000 | 0.141 | 0.525 | 0.581 | 1.000 | 1.000 | 0.938 | 0.132 | 0.138 | 1.000 | / |
|  | Avg Power | 1.000 | 0.393 | 0.675 | 0.733 | 0.997 | 0.994 | 0.946 | 0.411 | 0.384 | 1.000 | / |
|  | Type I Error | 0.077 | 0.067 | 0.074 | 0.076 | 0.057 | 0.074 | 0.932 | 0.062 | 0.123 | 0.789 | / |
|  | Avg Rank Sum | 21.00 | 107.37 | 46.17 | 40.72 | 21.19 | 21.45 | 136.82 | 101.84 | 108.26 | 26.13 | 35.63 |
| 0.5 | $X_1$ | 0.998 | 1.000 | 0.983 | 0.968 | 1.000 | 0.996 | 0.994 | 0.996 | 0.995 | 1.000 | / |
| | $X_{p_0+1}$ | 1.000 | 0.099 | 0.558 | 0.804 | 1.000 | 1.000 | 0.931 | 0.076 | 0.073 | 1.000 | / |
| | $X_{2p_0+1}$ | 1.000 | 0.082 | 0.703 | 0.864 | 1.000 | 0.983 | 0.946 | 0.076 | 0.154 | 0.998 | / |
| | $X_{3p_0+1}$ | 0.997 | 0.087 | 0.682 | 0.881 | 0.985 | 0.996 | 0.924 | 0.065 | 0.158 | 1.000 | / |
| | $Z_1$ | 0.996 | 0.997 | 0.798 | 0.930 | 0.995 | 0.981 | 0.969 | 0.996 | 1.000 | 0.999 | / |
| | $Z_2$ | 0.998 | 0.120 | 0.560 | 0.854 | 0.998 | 1.000 | 0.931 | 0.128 | 0.144 | 1.000 | / |
|  | Avg Power | 0.998 | 0.398 | 0.714 | 0.883 | 0.996 | 0.993 | 0.949 | 0.389 | 0.421 | 0.999 | / |
|  | Type I Error | 0.094 | 0.102 | 0.082 | 0.101 | 0.090 | 0.103 | 0.937 | 0.064 | 0.125 | 0.820 | / |
|  | Avg Rank Sum | 21.01 | 107.50 | 41.36 | 27.69 | 21.31 | 21.66 | 139.94 | 103.85 | 109.03 | 31.92 | 33.98 |

The true average rank sum of important features in Scenario 1 and 2 are 15 and 21, respectively. Feature importance identification performance comparison based on these three evaluation metrics are presented in Tables 1 and 2 while the prediction accuracy comparisons via *C*_*index*_ are displayed in Figure 1.

**Figure 1:**
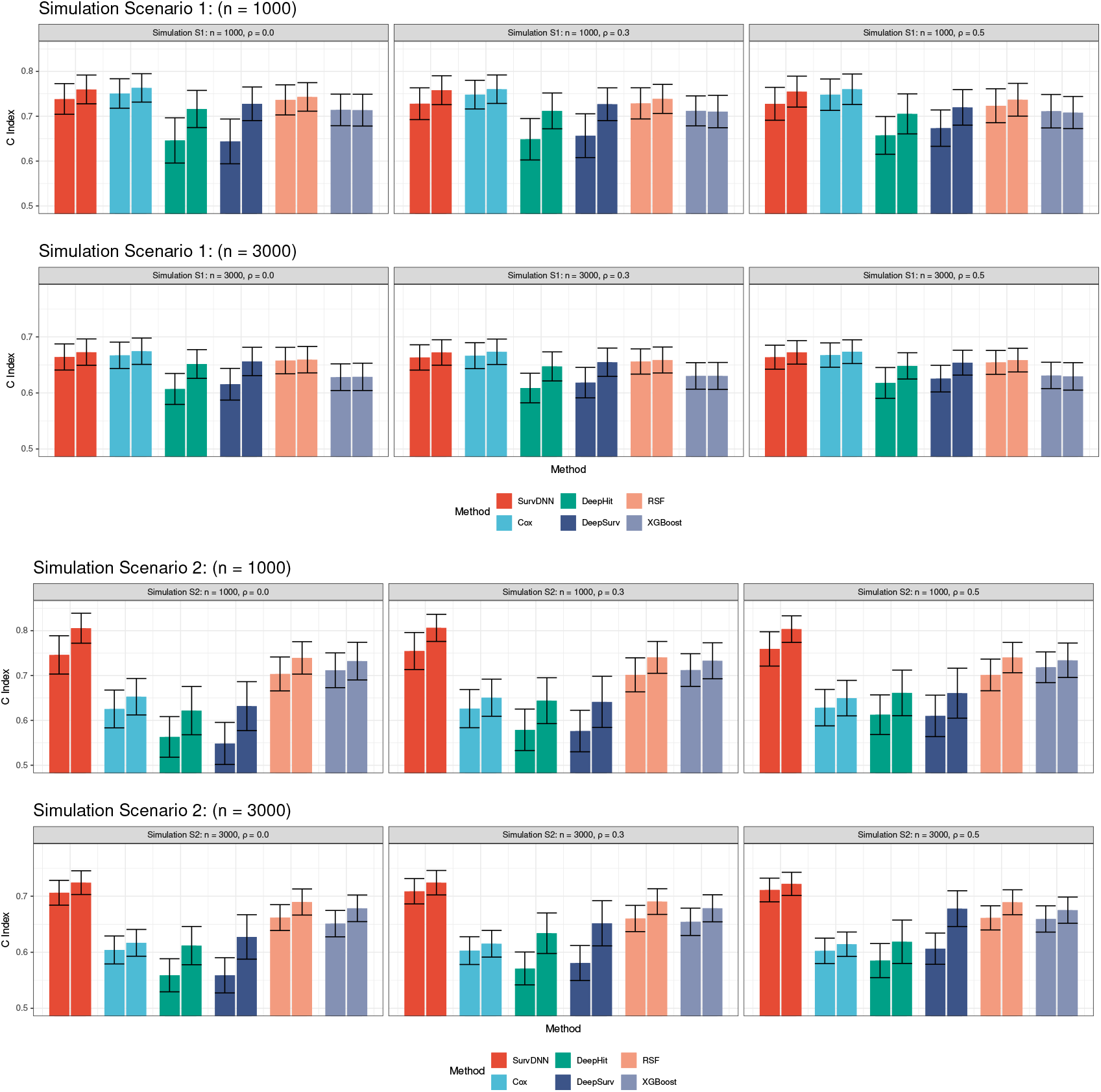
Survival Risk Prediction Comparison under Scenario 1 and 2.

In Scenario 1 with *n* = 1, 000, where the association between important features and survival outcome was linearly simulated, the Cox model showed superior performance in terms of power and average rank sum metrics compared to other methods due to the fact that Scenario 1 follows the assumption of the Cox model, leading to a favorable performance. However, due to limited sample size and potential overfitting, *p*-values from Cox models are slightly inflated. Tree-based methods like PermRSF and PermXGBoost demonstrated limited capability in identifying important features with weaker signals, such as *X*_1_ and 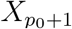. In contrast, our PermSurvDNN exhibited a stronger ability to detect features with weake signals compared to other machine-learning methods, preserved type I error at the nominal level, and performed slightly below the Cox model in power. An increase in *ρ* led to a slight inflation in the type I error rates for these permutation-based methods. However, our proposed PermSurvDNN still significantly outperformed PermRSF, PermXGBoost, and PermDeepSurv in terms of average rank sum and average power. Although Lasso-Cox and RSF performed well in selecting important features, it is important to note their highly inflated type I error rates due to their tendency to assign small yet significant importance scores to some null features. Wrapped with our proposed permutation-based framework, i.e., PermCox and PermRSF, the *p*−values for feature importance were better controlled. When *n* = 3, 000 and effect sizes were divided by 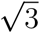, all methods showed improved ability in identifying features with weaker signals. We divided each effect size by 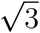 because leaving them unscaled would push statistical power close to 1 for all methods, making it impossible to compare their ability to detect weaker signals. The comparison results among different methods remained consistent with those in Scenario 1 with *n* = 1, 000.

In Scenario 2 with *n* = 1000, the permutation test based on machine learning models exhibited more compelling power than the Cox model. PermSurvDNN, PermRSF, and PermXGBoost effectively identified the quadratic term 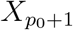, with PermSurvDNN also excelling in detecting the interaction between 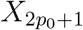 and 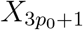. Although PermSurvDNN’s type I error rate increased with rising *ρ*, it consistently outperformed other competing methods in the average rank sum metric, affirming its effectiveness in important feature identification and its utility in real-world studies. When *n* increased to 3000 and effect sizes were divided by 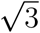, PermSurvDNN demonstrated a strong capacity for identifying features with smaller coefficients. Its corresponding average rank sum metric was even closer to the oracle value of 21.

Regarding survival risk prediction, we observed that the prediction accuracy of each survival model generally improved following the selection of important features, e.g. SurvDNN versus PermSurvDNN, and DeepSurv versus PermDeepSurv. SurvDNN’s prediction performance was comparable to the Cox model in linear Scenario 1 and surpassed all other models in Scenario 2 in terms of the C-index. The evaluation of prediction accuracy using the integrated Brier score yielded similar results, as shown in Appendix Section 1.1. This superior performance can be attributed to SurvDNN’s ability to capture complex associations for accurate and robust outcome predictions, and the permutation procedure that effectively identifies important features. Furthermore, DeepSurv consistently underperformed compared to SurvDNN in all scenarios for both feature selection and outcome prediction. This inferior performance can be attributed to DeepSurv’s lack of the bagging and filtering DNN algorithm proposed in this paper. Another deep-learning-based method, DeepHit, initially performed even worse when using default hyper-parameters. However, after tuning additional hyper-parameters, its performance became comparable to Deep-Surv, though it still remained inferior to the proposed SurvDNN. This may be due to the limitations of using traditional DNNs in DeepHit.

In summary, across both linear and non-linear simulation scenarios, PermSurvDNN achieved superior power and tighter feature ranking (lower average rank sum) while controlling type I error, compared to PermCox, PermRSF, and other machine-learning methods (Figure 1; Tables 1 and 2). Notably, under moderate correlation (*ρ* = 0.3 ∼ 0.5), PermSurvDNN uniquely maintained high power for identifying weak predictors 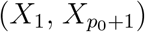 where tree-based methods faltered. These results underscore PermSurvDNN’s robust ability to detect important features associated with survival outcomes, regardless of the complexity of the predictor–outcome relationships.

### 2.2 Real Application

To evaluate the effectiveness of our approach in clinically realistic settings, we applied PermSurvDNN and competing models to two large, real-world cohorts: the Surveillance, Epidemiology, and End Results (SEER) registry and the Molecular Taxonomy of Breast Cancer International Consortium (METABRIC) study, with the former representing a low-dimensional setting and the latter a high-dimensional scenario. In both applications, PermSurvDNN not only robustly and consistently identified clinically validated critical biomarkers and risk factors that other methods frequently missed but also achieved superior predictive accuracy.

#### 2.2.1 Surveillance, Epidemiology, and End Results (SEER) Study

In this low-dimensional setting, we analyzed 4,668 hepatocellular-carcinoma (HCC) patients (2018–2021) from SEER (denoted as SEER-HCC), using 19 covariates spanning demographics, tumor characteristics, biomarkers, and treatments. After an 80/20 train/test split, each model was trained on the training set, and PermFIT was used to select important features (with p-value ≤ 0.05 as cut-off). This process was replicated for 100 times.

As shown in Figure 2a, PermSurvDNN consistently identified 12 key predictors including alpha-fetoprotein (AFP), liver fibrosis score, age, tumor size, grade, and others - each of which has been clinically validated in HCC prognosis [29, 30]. In contrast, PermRSF and other competing methods, such as PermCox, frequently omitted age and fibrosis score despite their established prognostic importance. Notably, patient age has been demonstrated as a prognostic determinant in a large meta-analysis study [36]. However, as shown in Figure 2a, PermSurvDNN is the only method that consistently identifies age as a significant feature associated with HCC patients’ survival outcomes. This observation also raises concerns about the reliability of other competing methods in identifying important features and biomarkers associated with complex disease survival outcomes. Alpha-fetoprotein and liver fibrosis reflect tumor biology and underlying liver health, respectively. Their robust selection underscores PermSurvDNN’s ability to uncover both molecular and clinical predictors. SurvDNN achieved the highest C-index on both full-feature and reduced-feature models, i.e., PermSurvDNN yielded a test C-index of 0.785, outperforming PermCox (0.780), PermRSF (0.774), PermDeepSurv (0.774), PermDeepHit (0.771), and PermXGBoost (0.756) (Figure 2b). This consistent performance even after feature selection underscores PermSurvDNN’s ability to robustly deliver reliable and clinically meaningful risk factor as well as biomarker identification.

**Figure 2:**
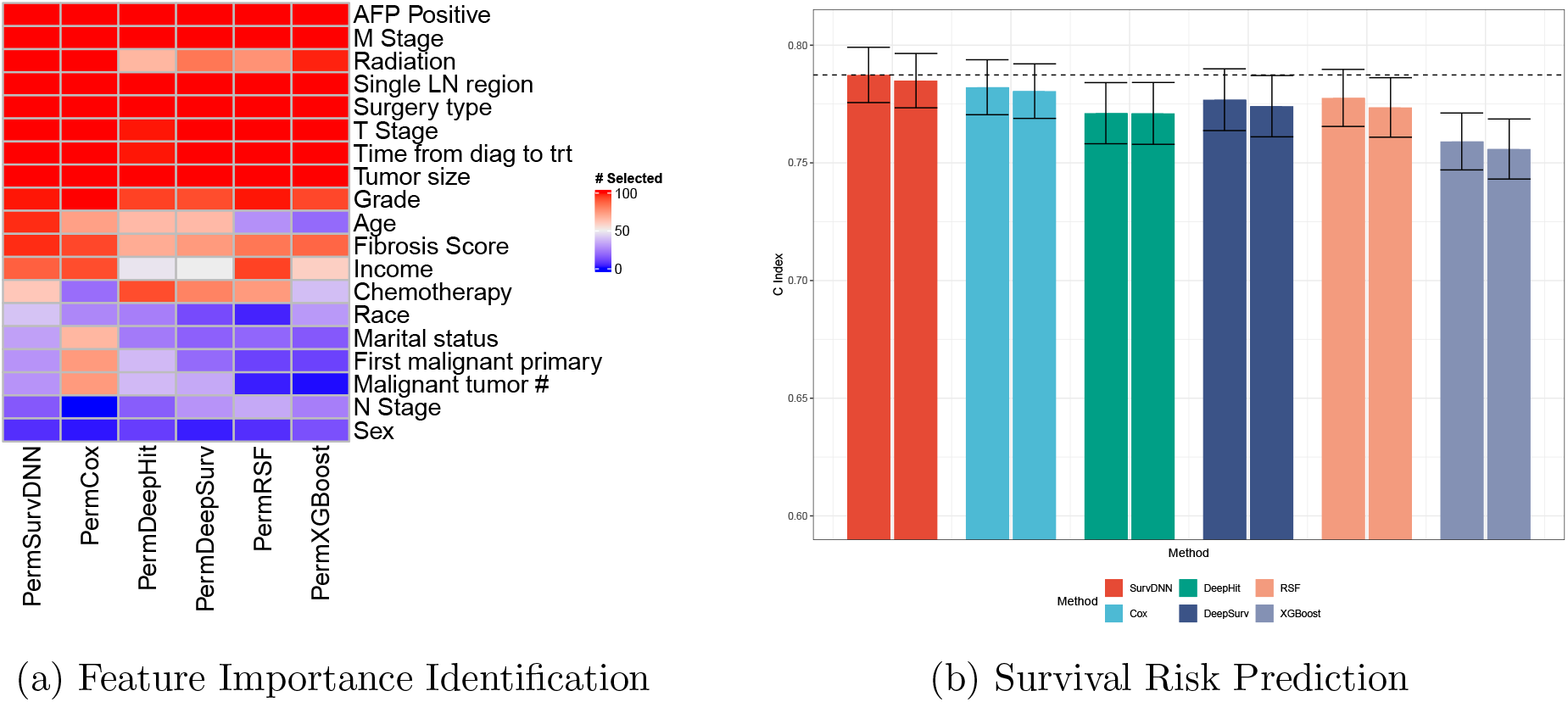
Model Performance Comparison for SEER-HCC Study.

To further demonstrate the superiority of PermSurvDNN in identifying important and clinically meaningful biomarkers for complex disease survival outcomes as well as its capability to accurately predict risk scores, we present Kaplan–Meier curves based on testing samples stratified by the identified biomarkers (e.g., AFP and fibrosis score) and the predicted risk scores from PermSurvDNN. These scores are dichotomized at the median into high-and low-risk groups, as shown in Figure 3. As illustrated in this figure, patients’ survival statuses are well separated by both the identified biomarkers and the predicted risk scores. This clearly highlights the practical clinical utility of PermSurvDNN in determining features and biomarkers associated with complex disease survival outcomes, where other competing methods often fail to perform effectively. Most importantly, reliably identifying key risk features and biomarkers for complex diseases is critical for gaining insights into disease mechanisms.

**Figure 3:**
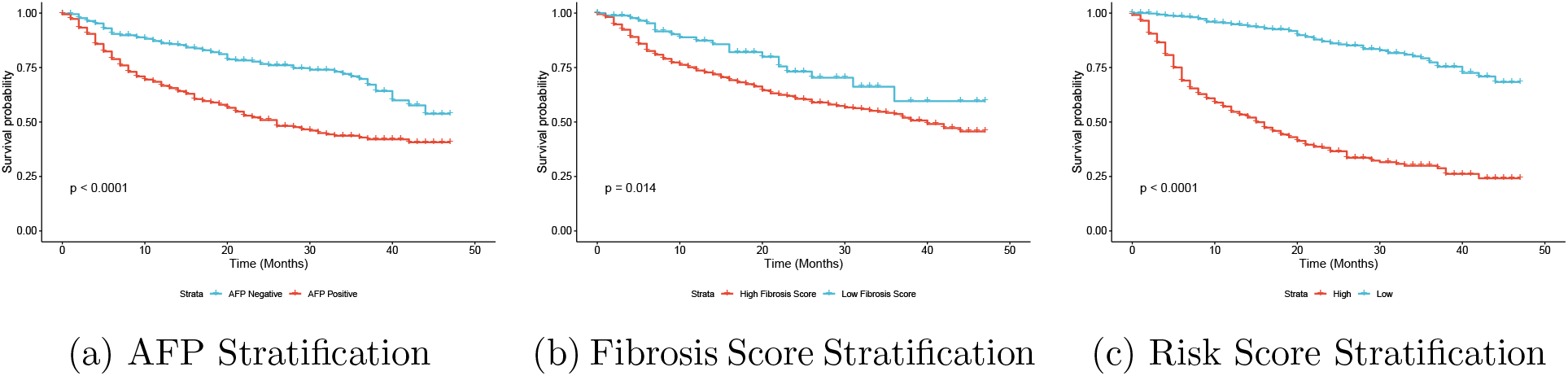
Kaplan–Meier Curve Stratification in SEER-HCC Study.

#### 2.2.2 Molecular Taxonomy of Breast Cancer International Consortium Study

To evaluate the performance of PermSurvDNN in high-dimensional settings, we applied it to the Molecular Taxonomy of Breast Cancer International Consortium (METABRIC) dataset. From the METABRIC cohort, we retained 1,684 patients after filtering for missing data and included 67 features comprising 20 clinical variables and 47 mutations by excluding mutations with a frequency *<* 100 from the original set of 172 mutations. Using the same 80/20 train-test split, PermSurvDNN consistently selected ten significant features (Figure 4a), including TP53 mutation, tumor size, Nottingham Prognostic Index (NPI), and BRCA subtype. These features have been validated in previous studies as critical factors associated with breast cancer survival outcomes. For instance, the TP53 gene, a key tumor suppressor, was consistently selected by PermSurvDNN and PermRSF. Prior research has linked TP53 mutations to poorer survival outcomes in breast cancer [37, 38]. Larger tumor size has consistently been linked to worse survival outcomes [39, 40]. Similarly, the NPI, a metric used to predict survival in operable primary breast cancer [41], has been widely recognized as an important prognostic factor [42]. The BRCA subtype has also been confirmed as a crucial survival-related feature in prior clinical studies [43]. In contrast, PermCox and Lasso-Cox failed to reliably recover the BRCA subtype. PermSurvDNN uniquely captured both NPI and BRCA subtype across all folds, highlighting SurvDNN’s strength in modeling complex associations. Predictive results (Figure 4b) show that both SurvDNN and PermSurvDNN outperform all competing algorithms, confirming the method’s superior accuracy and interpretability in a real-world breast cancer cohort.

**Figure 4:**
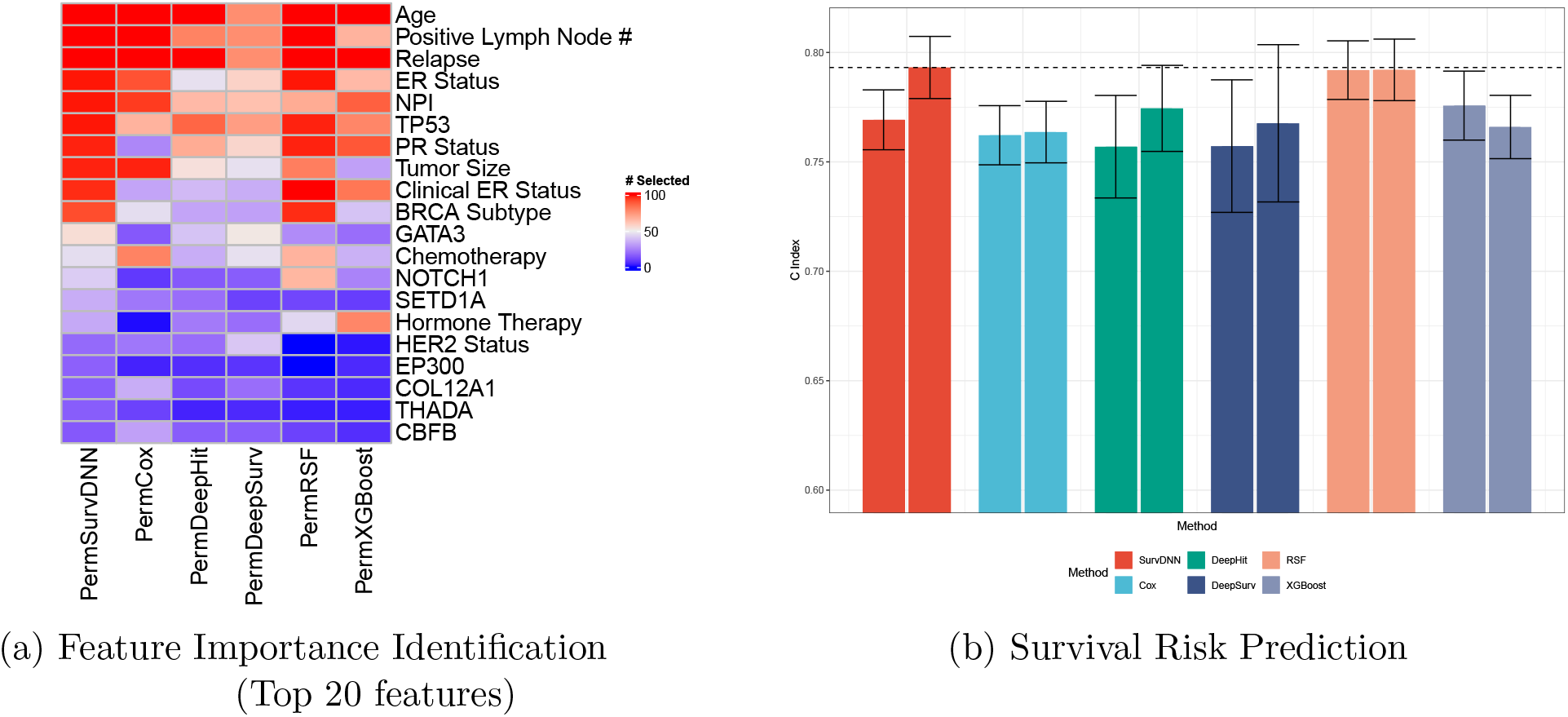
Model Performance Comparison for METABRIC Study.

As additional evidence of PermSurvDNN’s advantage in recovering clinically meaningful prognostic factors and producing accurate risk stratification, we show Kaplan–Meier curves from a representative METABRIC split, stratified by identified factors (e.g., tumor size, NPI) and by PermSurvDNN-predicted risk scores (Figure 5). Both factor-based and model-based strata yield clear survival separation, indicating strong out-of-sample discrimination. Notably, while tumor size and NPI emerge as significant in this analysis - consistent with prior studies [39, 40], competing methods often show reduced power to detect these variables or miss them entirely, underscoring PermSurvDNN’s ability to surface clinically salient signals with direct relevance for risk stratification in practice.

**Figure 5:**
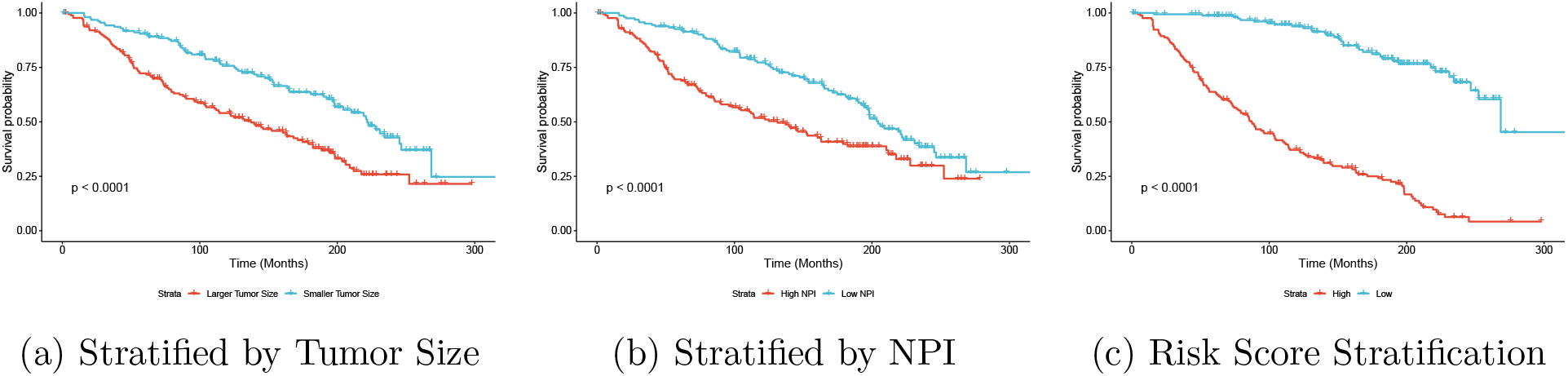
Kaplan–Meier Curve Stratification in METABRIC Study.

## 3 Discussion

In this paper, we introduce SurvDNN, an enhanced deep learning framework for survival outcome modeling. SurvDNN builds on the conventional Cox-based DNN by incorporating a bootstrapping-based regularization strategy to mitigate overfitting and a stability-driven filtering algorithm to improve model robustness. We extend it with the Permutation-based Feature Importance Test (PermFIT) for survival settings, yielding PermSurvDNN to rigorously quantify individual biomarker contributions and deliver reliable risk predictions. This approach enables us to accurately identify important biomarkers and interpret each feature’s contribution within high-dimensional, non-linear, non-additive contexts for survival outcomes. Through extensive simulations and applications to real-world complex disease datasets, SurvDNN consistently outperforms existing machine learning methods in both predictive accuracy and biomarker discovery. An open-source R package implementing SurvDNN is publicly available on GitHub, facilitating its adoption for precision-medicine research. Our methods achieve superior performance across both simple parametric settings and complex survival scenarios where standard linear-additive survival models fail. Overall, our findings confirm that PermSurvDNN combining bootstrapped regularization, stability filtering, and PermFIT offers a powerful, robust, interpretable tool for biomarker-driven survival modeling regardless of whether the relationship between predictive features and survival outcomes is simple or complex.

It is important to note that permuting one feature at a time can alter the predictors’ covariance structure. Conditional permutation tests could mitigate this issue, though they introduce additional assumptions and are beyond this paper’s scope. Furthermore, our proposed method is specifically designed for right-censored, single-event survival data; extending it to interval-censored or competing-risk analyses warrants future work. Additionally, it’s worth noting that the proposed DNN method and permutation-based test entail a more intensive computational workload. We intend to focus on addressing these limitations, including optimizing computational efficiency and exploring advanced permutation schemes, in our future research.

## 4 Methods

### 4.1 Enhanced Deep Neural Network for Complex Survival Data

Before presenting the details of SurvDNN, we introduce some notations. Assuming that the training data has *n* subjects, where the survival outcome is only subject to right censoring, we denote the *p* dimensional feature measurements as ***X*** = (***x***_**1·**_, · · ·, ***x***_***n*·**_)*′* = (***x***_**·1**_, · · ·, ***x***_**·*p***_) with ***x***_***i*·**_ = (*x*_*i*1_, *x*_*i*2_, · · ·, *x*_*ip*_)*′* and ***x***_**·*j***_ = (*x*_1*j*_, · · ·, *x*_*nj*_)*′, i* ∈{1, · · ·, *n* }, *j* ∈ {1, · · ·, *p*} referring the associated feature vector of individual *i* and feature *j*, respectively. For each subject *i*, the observed event time is given by *T*_*i*_ = min(*T*_true,*i*_, *C*_*i*_), where *T*_true,*i*_ represents the true underlying event time, and *C*_*i*_ is the follow-up time. The event indicator *δ*_*i*_ is defined as *δ*_*i*_ = 𝕀 (*T*_true,*i*_ ≤ *C*_*i*_), where *δ*_*i*_ = 1 if the event occurred, or uncensored, and *δ*_*i*_ = 0 if the event is right-censored. In summary, we have training data 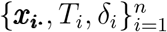.

We follow the Cox proportional hazard model and assume *h*(*t*|***x***_***i*·**_) = *h*_0_(*t*) exp[*ξ*(***x***_***i*·**_|***θ***)], where *h*(*t* |***x***_***i*·**_) is the conditional hazard function of survival time, *h*_0_(*t*) is the unspecified baseline hazard function, and ***θ*** is the collection of model parameters. The partial log-likelihood of the model is therefore

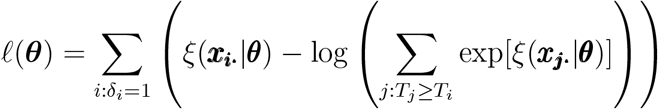

Instead of modeling *ξ* parametrically as in the traditional Cox proportional hazard where *ξ*(***x***_***i*·**_|***θ***):= ***x***_***i*·**_*′* ***β***, SurvDNN models this function flexibly using a *K*-hidden layer DNN. That is, each hidden layer *k* ∈ {1,…, *K*} produces an intermediate representation, ***η***^(*k*)^ through an activation function ***g***_*k*_. Specifically, ***η***^(*k*)^ = ***g***_*k*_[***b***^(*k*)^ +***W*** ^(*k*)^***η***^(*k*−1)^], where *n*_(*k*)_ is the dimension of ***η***^(*k*)^, ***b***^(*k*)^ is the bias vector with a length of *n*_(*k*)_ and ***W*** ^(*k*)^ is an *n*_(*k*)_ *× n*_(*k*−1)_ weight matrix. Function ***g***_*k*_ applies an activation function *g*_*k*_ element-wise to the *n*_(*k*)_ dimensional vector ***b***^(*k*)^ + ***W*** ^(*k*)^***η***^(*k*−1)^. We take the common practice approach and use ReLU (rectified linear unit function) for all *g*_*k*_’s (*k* = 1, · · ·, *K*) [44]. For the first hidden layer, ***η***^(0)^ is simply *n* elements of the original *p*-dimensional feature matrix ***X***. Finally, the *K*^*th*^ hidden layer sets *ξ*(·|***θ***) = ***b***^(*K*+1)^ + ***W*** ^(*K*+1)^***η***^(*K*)^. In summary, SurvDNN takes ***x***_***i*·**_ as an input with 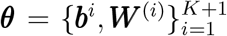 being the collection of parameters. To ensure the identifiability of *ξ*, we add a constraint to the last layer bias ***b***^(*K*+1)^ such that 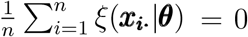. In addition, to deal with the high-dimensionality of covariates, we further introduce an ℓ_1_ penalty to the DNN loss function on the weight matrices ***W*** ^(*k*)^, (*k* = 1, · · ·, *K* + 1). Eventually, 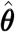, the estimate of ***θ***, is obtained by minimizing the following risk function:

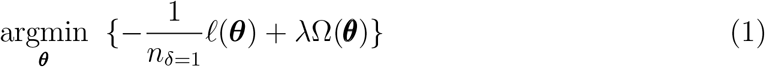

where *λ* is a penalty weight, 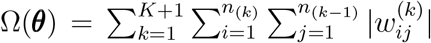, with 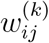 being the *ij*^th^ element in the weight matrix 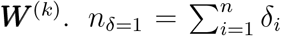 denotes number of observations who are not censored. We optimize (1) by using the mini-batch stochastic gradient descent algorithm [45, 46], along with adaptive moment estimation (Adam) [47]. Let the relative risk prediction of subject *i* with feature profile ***x***_***i*·**_ be 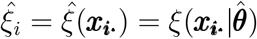.

Under finite sample settings, DNN models may not be stable. To overcome this issue, we adopt a bootstrap aggregating and filtering procedure as introduced by Mi et al. [16]. First, we generate bootstrap samples by randomly sampling the training set with replacement *B* times. For each bootstrapped sample, we fit a DNN model, reserving the out-of-bag (OOB) samples as a validation set. Denote the corresponding estimated relative risk prediction 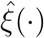 from the *b*^*th*^ bootstrapped samples as 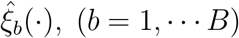. Traditionally, these estimates are averaged to get the final estimate of *ξ* as 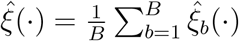. However, due to the random parameter initialization, some of these bagged DNN models may not approximate *ξ* well. Research has suggested using a subset of the bagged models that fit the data well [16, 48]. To exclude poorly performing models from the final ensemble, we adopt a performance score, *v*_*b*_, for the *b*^*th*^ bootstrapped sample as:

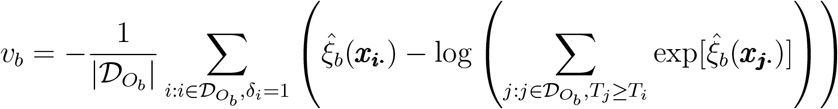

where 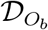 is the OOB samples with an associated sample size 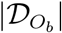.

To determine the optimal subset of DNNs for the final ensemble, we first rank the DNNs based on their performance scores, denoting as *v*_(1)_ ≥ · · · ≥ *v*_(*B*)_ with the corresponding function estimate as 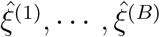. By aggregating the top *q* DNNs, the new aggregated prediction becomes

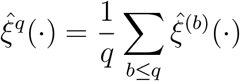

The optimal number of DNNs utilized by the ensemble, *q*_opt_, is determined by minimizing the training loss such that:

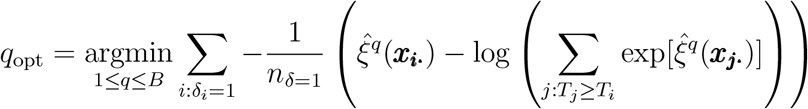

SurvDNN utilizes the revised bagging prediction 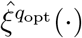 as the final prediction.

To measure the prediction accuracy in survival analysis, we use Harrell’s C-index, which is a generalization of the area under the receiver operating characteristic (ROC) curve (AUC) for censored data [49] as follows:

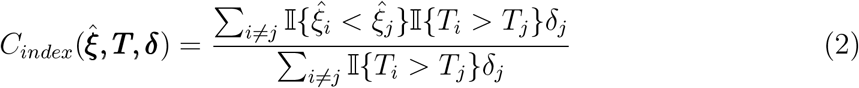

where 𝕀 (·) is an indicator function,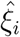 is the relative risk prediction of subject *i*. The higher the *C*_*index*_, the better the relative risk prediction.

We further utilize the integrated Brier score (IBS) to measure the prediction performance. Following Kvamme et al. [50], we have the Brier score (BS) at a given time *t* as follows:

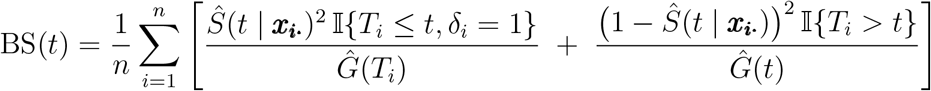

where *Ŝ*(*t* | ***x***_***i*·**_) is the survival probability prediction of the model to be evaluated, *Ĝ*(*t*) is the Kaplan-Meier estimate of the censoring survival function, *P* (*C*_*i*_ *> t*), and it is assumed that the censoring times and survival times are independent.

The integrated Brier score (IBS) is defined as:

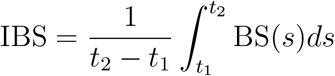

In practice, we approximate this integral by applying numerical integration (i.e., trapezoidal rule), and we let the time span be the duration of the test set.

### 4.2 Robust Feature Importance Identification for Complex Survival Data via Machine Learning Models

In traditional machine learning models with non-survival data, to assess the importance of a given feature, PermFIT [19] empirically quantifies the difference in prediction accuracy between using its observed values and using permuted values as input. In essence, it assesses how much the predictive performance of a model changes when the actual feature values are replaced with randomly permuted ones, based on which the importance of the feature can be evaluated. Following Mi et al. [19], we extend PermFIT to survival models as follows: firstly, with a trained survival model 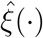, and error measure 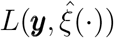 which is defined later, we can estimate the prediction error with observed feature 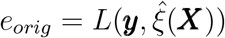; secondly, for each feature vector ***x***_**·*j***_, *j* ∈ {1, · · ·, *p*}, we can then generate feature matrix ***X***_*perm,j*_ by permuting feature vector ***x***_**·*j***_ without replacement, denoted as ***x***_**·*j***_^Perm^, resulting in ***X***_*perm,j*_ = (***x***_**·1**_, · · ·, ***x***_**·(*j*−1)**_, ***x***_**·*j***_^Perm^, ***x***_**·(*j*+1)**_, · · · ***x***_**·*p***_). Define the prediction error associated with 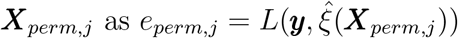 and the permutation-based feature importance for *j*^*th*^ feature Δ_*j*_ = *e*_*perm,j*_ − *e*_*orig*_. By adopting C-index as the prediction performance measurement of survival outcomes, we define the error measure 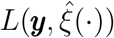 as 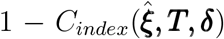. The feature importance score for the *j*^th^ feature is estimated as 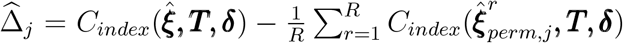, where *R* is the total number of random permutations, and 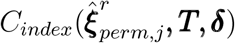 is calculated similarly to the original C-index with equation (2) except that the relative risk prediction 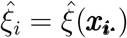 is replaced by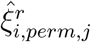which is the relative risk prediction based on the *i*^*th*^ row of the permuted feature matrix ***X***_*perm,j*_ in the *r*^*th*^ permutation.

To avoid the potential overfitting of different machine learning algorithms and increase the power of important feature identification, we further employ *B*-fold cross-fitting strategy [19]. Here, we randomly divide the data into *B* folds and denote the corresponding data as *V*_1_, · · ·, *V*_*B*_. For each of *V*_*b*_ (*b* = 1, · · ·, *B*), 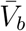 denotes the complementary set of *V*_*b*_, which is used to fit the survival model 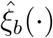, and |*V*_*b*_| = the cardinality of *V*_*b*_. Let 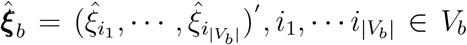 be the survival prediction vector of the data in validation set *V*_*b*_ given by corresponding fitted model 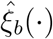, and 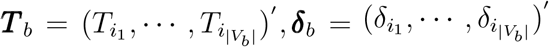 be the time and event status vector of the data in validation set *V*_*b*_. Then, we can permute the *j*^*th*^ feature of the data in validation set *V*_*b*_, and get the estimated feature importance score of the *j*^th^ feature 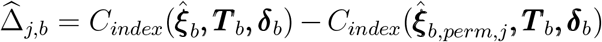.To deal with the randomness of each permutation, we repeat this process by *R* times, get *R* estimated feature importance score 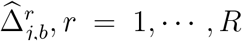in each permutation and each fold, and then average to get the final estimate: 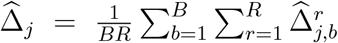 and 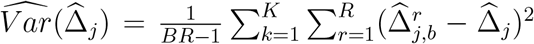. We hereby summarize our permutation-based feature importance test for survival analysis in Algorithm 1.

Similarly, we can apply this permutation procedure to various survival machine learning models to evaluate feature importance. We denote the feature importance computed by a given model as PermModel - for example, PermCox and PermSurvDNN for the Cox model and SurvDNN, respectively. The important features identified by PermFIT can then be used to refine each model. We also use PermModel to refer to the refined model’s survival prediction performance - for instance, PermSurvDNN denotes the prediction performance of the SurvDNN model when using the selected important features. Detailed descriptions of the survival prediction procedures for each algorithm, including RSF, XGBoost, DeepHit, and DeepSurv, are provided in the Appendix.

#### Algorithm 1 Permutation Feature Importance Test for SurvDNN

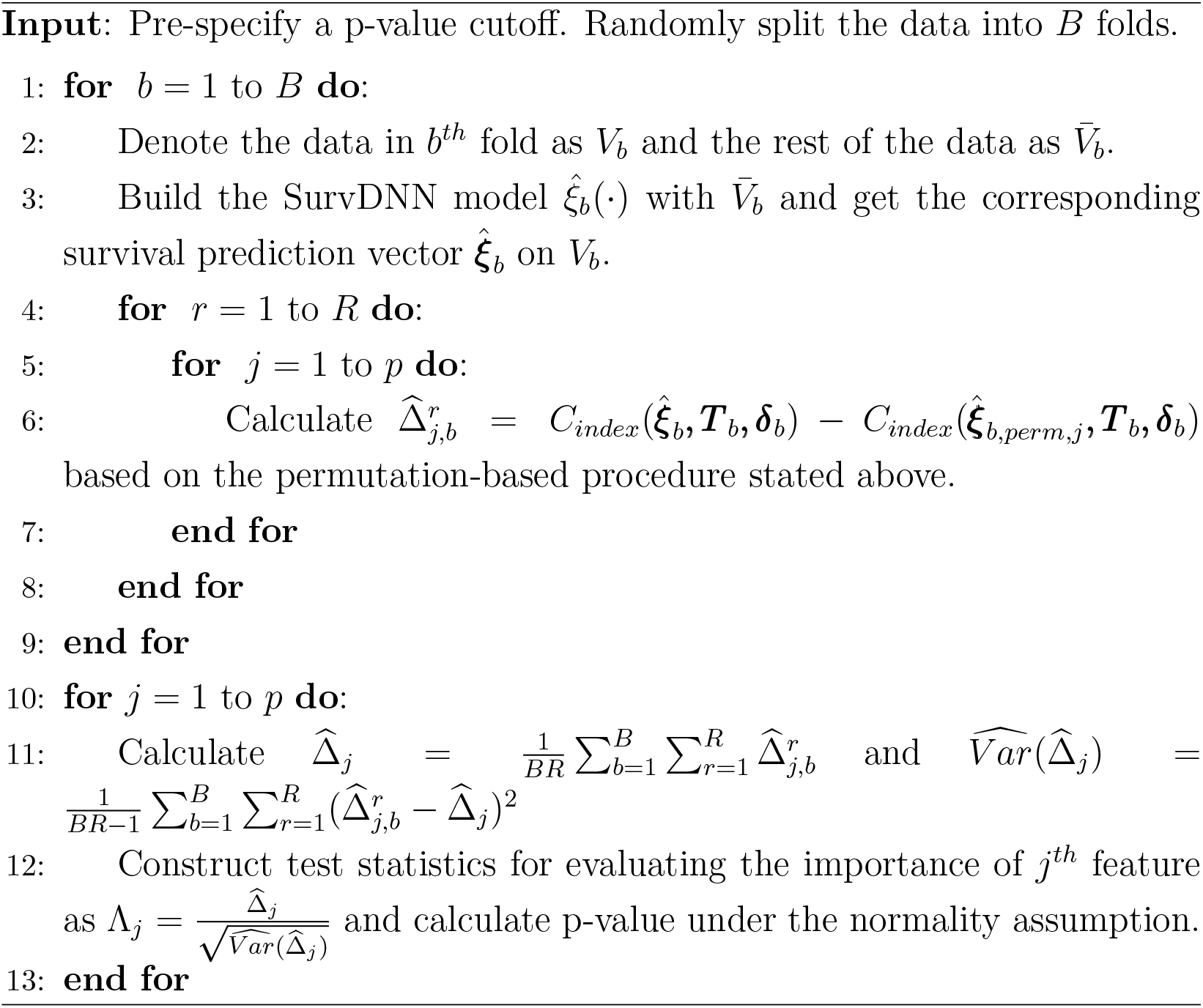

## Acknowledgement

This work was partially supported by the National Institute of Health Grants R01LM014407 and 1R01HL173044.

## Author contributions

S.W. implemented the algorithms in “SurvDNN” for the proposed method and performed numerical analyses. All authors contributed to the methodology development and writing the manuscript.

## Data availability

We utilize Surveillance Research Program, National Cancer Institute SEER*Stat software (seer.cancer.gov/seerstat) version 8.4.5, to extract the SEER-HCC dataset from Surveillance, Epidemiology, and End Results (SEER) Program (www.seer.cancer.gov) SEER*Stat Database: Incidence - SEER Research Data, 17 Registries, Nov 2023 Sub (2000-2021) - Linked To County Attributes - Time Dependent (1990-2022) Income/Rurality, 1969-2022 Counties. (https://seer.cancer.gov/data-software/documentation/seerstat/nov2023/). The METABRIC dataset was downloaded from cBioPortal (https://www.cbioportal.org/study/summary?id=brca_metabric).

## Code availability

An R package implementing the proposed framework is available on GitHub (https://github.com/BZou-lab/SurvDNN).

## SUPPLEMENTARY MATERIAL

**Appendix.pdf** includes the implementation details of existing machine learning survival models, such as the R packages used and the tuning of hyperparameters, and also the prediction performances measured by integrated Brier score.

