## Appendix for "An Interpretable Deep Learning Framework for Biomarker Discovery in Complex Disease Survival Outcomes"

#### 1 Performance Comparison via Integrated Brier Score

In addition to using the C-index to evaluate model performance for survival outcomes, we also compare each model's performance across various simulation settings and real-data applications using the integrated Brier score. Supplementary Figures 1 and 2 present predictive performance - measured by the integrated Brier score - in simulation scenarios 1 and 2, while supplementary Figures 3 and 4 show the integrated Brier scores for the SEER-HCC and METABRIC studies.

##### 1.1 Simluation Studies

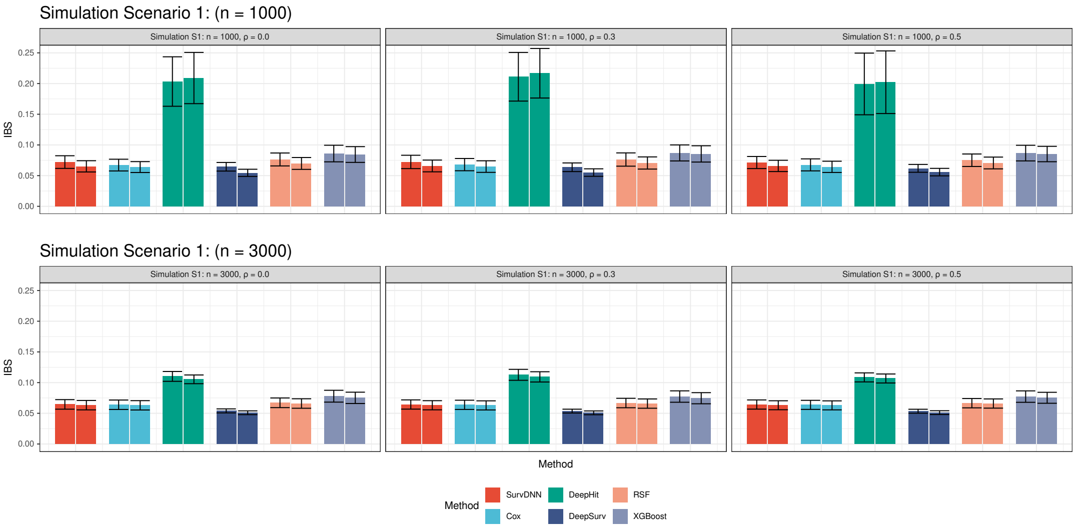


Figure 1: Survival Risk Prediction Comparison under Scenario 1

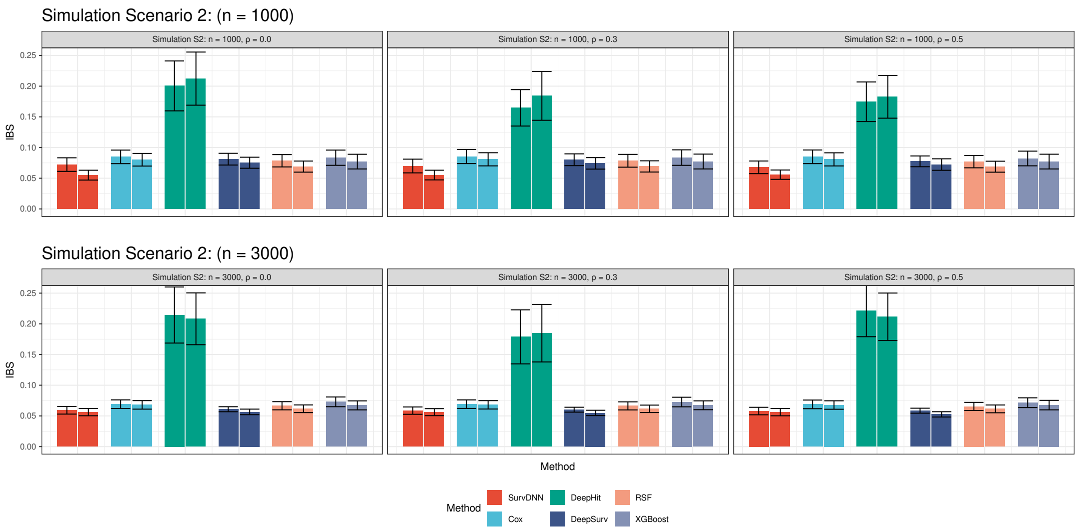


Figure 2: Survival Risk Prediction Comparison under Scenario 2

#### 1.2 SEER-HCC

Supplementary Figure 3 presents the evaluation of prediction performance based on the integrated Brier score in SEER-HCC study.

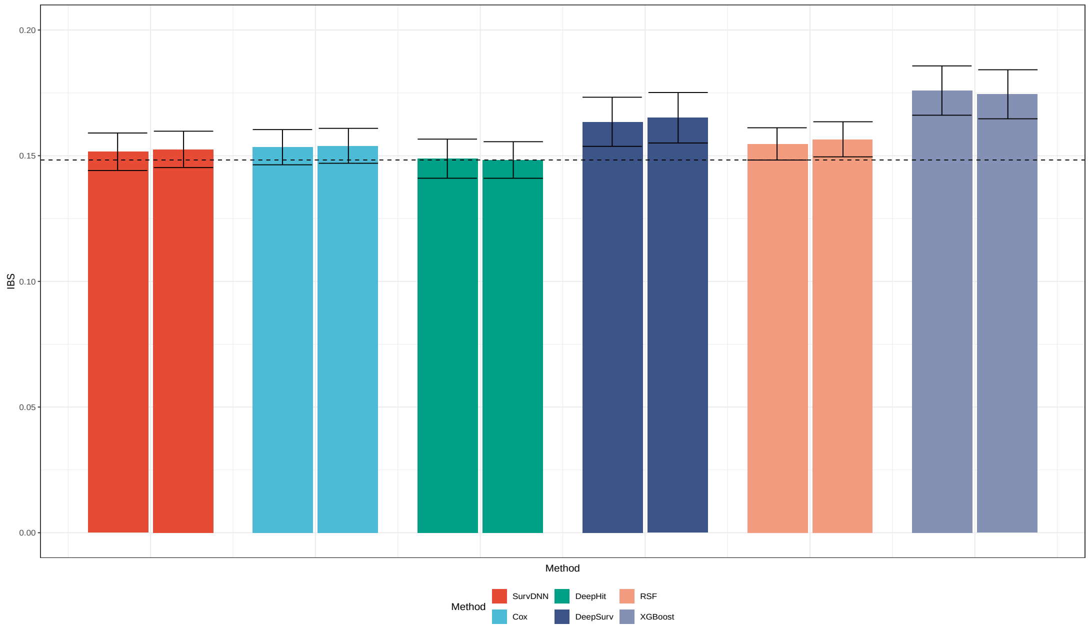


Figure 3: Survival Risk Prediction Comparison for SEER-HCC Study

#### 1.3 METABRIC

Supplementary Figure 4 presents the evaluation of prediction performance based on the integrated Brier score in METABRIC study.

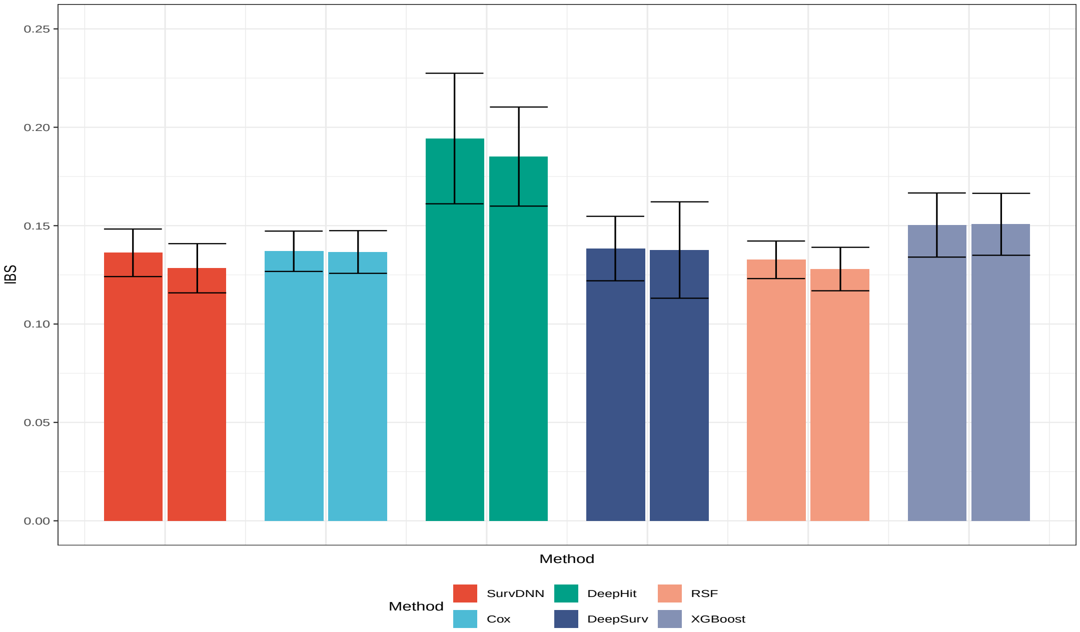


Figure 4: Survival Risk Prediction Comparison for METABRIC Study

### 2 Description of Existing Machine Learning Survival Models Implementation Details

##### Cox Proportional Hazard Model

The Cox proportional hazard (CPH) model assumes the functional relationship of the hazard to be $h(t|\boldsymbol{X_{i\cdot}}) = h_0(t)\exp(\boldsymbol{X_{i\cdot}}'\boldsymbol{\beta})$. Consequently, the relative risk prediction of the CPH model is defined as $\hat{\xi}_{i,\text{Cox}}(\boldsymbol{X_{i\cdot}}) = \exp(\boldsymbol{X_{i\cdot}}'\hat{\boldsymbol{\beta}})$. CPH model is implemented via the R package "survival" [1].

#### Lasso - Cox

The model assumption of Lasso-Cox is the same as CPH, but the regression coefficient $\boldsymbol{\beta}$ is estimated by minimizing the negative partial likelihood plus an $l_1$ penalty. The relative risk prediction of Lasso-Cox model is $\hat{\xi}_{i,\text{LASSO-Cox}}(\boldsymbol{X_{i\cdot}}) = \exp(\boldsymbol{X_{i\cdot}}'\hat{\boldsymbol{\beta}}_{\textbf{Lasso}})$. The Lasso-Cox model is implemented via R package "glmnet" [2]. The penalty weight is determined by using function *cv.glmnet* implemented in "glmnet".

#### Random Survival Forest (RSF)

Random Survival Forests (RSF) involve drawing B bootstrap samples, randomly selecting a subset of features, growing a survival tree on each bootstrap sample using log-rank test statistic-based splitting rules, computing the cumulative hazard function (CHF) for each tree, and ultimately averaging them to derive the ensemble CHF. The survival prediction given by RSF is the expected mortality: $M(\boldsymbol{X_{i\cdot}}) = \sum_{j=1}^{m} \hat{H}_h(t_j|\boldsymbol{X_{i\cdot}})$, where the test data $\boldsymbol{X_{i\cdot}}$ is dropped down the trained survival forest and reaches the terminal node $h$, $\hat{H}_h(t|\cdot)$ is the Nelson-Aalen estimate of the CHF for terminal node $h$ at time $t$, and $(t_1, \cdots, t_m)$ are the entire set of unique event times for the training data. Detailed description of such expected mortality can be found elsewhere [3]. We denote the relative risk prediction of RSF as $\hat{\xi}_{i,\text{RSF}} = M(\boldsymbol{X_{i\cdot}})$. The RSF model is implemented via R package "randomForestSRC" [4]. For all the analyses, we used *ntrees = 500* and set other tuning parameters as default.

#### Extreme Gradient Boosting (XGboost)

XGBoost is a gradient-boosting ensemble model consisting of a group of decision trees, which can deal with survival data by learning each tree using the survival loss function setting. The model assumption of the XGboost is $h(t|\boldsymbol{X_{i\cdot}}) = h_0(t)\exp[\xi(\boldsymbol{X_{i\cdot}})]$, and $\hat{\mathcal{T}}(\boldsymbol{X_{i\cdot}})$ represents the output from a decision tree ensemble to approximate $\xi(\boldsymbol{X_{i\cdot}})$, given input

$\boldsymbol{X_{i\cdot}}$. The survival risk prediction of XGboost is denoted as $\hat{\xi}_{i,\text{XGboost}} = \hat{\mathcal{T}}(\boldsymbol{X_{i\cdot}})$. The XGboost model is implemented via R package "xgboost" [5]. For all the analyses, we used task parameter *objective = survival:cox*, *nrounds = 50* and set other tuning parameters as default.

**DeepSurv**

DeepSurv is a deep neural network with model assumption: $h(t|\boldsymbol{X_{i\cdot}}) = h_0(t)\exp[\xi(\boldsymbol{X_{i\cdot}})]$. The network is trained by setting the loss function to be the average negative log partial likelihood plus a $\ell_2$ panalty, and the output of the network $\hat{\xi}(\boldsymbol{X_{i\cdot}})$ is trained to approximate $\xi(\boldsymbol{X_{i\cdot}})$. DeepSurv could estimate the survival probability of patient $i$ at time $t_j$: $\hat{S}(t_j|\boldsymbol{X_{i\cdot}})$. Based on the concept of expected mortality [6] and the requirement of the permutation feature importance test, we define the survival risk prediction of DeepSurv to be $\hat{\xi}_{i,\text{DeepSurv}} = \sum_{j=1}^{m} -\log[\hat{S}(t_j|\boldsymbol{X_{i\cdot}})]$, where $(t_1, \cdots, t_m)$ are the entire set of unique event times for the training data. The DeepSurv model is implemented via R package "survivalmodels" [7]. For all the analyses, we kept the hyperparameters to be the same as we used previously [8], i.e., the number of hidden layers to be 4 with 50, 40, 30, 20 hidden nodes at each layer, batch size to be 50, epochs to be 1000 with early stopping.

**DeepHit**

DeepHit uses a multi-task deep neural network to learn the distribution of survival times directly. It uses a shared sub-network and cause-specific sub-networks to estimate $\hat{P}(t,k|\boldsymbol{X_{i\cdot}})$ that the patient will experience the event $k$ at time $t$. DeepHit could also estimate the survival probability of patient $i$ at time $t_j : \hat{S}(t_j, 1|\boldsymbol{X_{i\cdot}}) = 1 - \hat{P}(t_j, 1|\boldsymbol{X_{i\cdot}})$. Similar with above, we define the survival risk prediction of DeepHit as $\hat{\xi}_{i,\text{DeepHit}} = \sum_{j=1}^{m} -\log[\hat{S}(t_j, 1|\boldsymbol{X_{i\cdot}})]$. The DeepHit model is implemented via R package "survivalmodels" [7]. We used the same hyperparameters of network structure and training step described above.

Further, DeepHit is a discrete survival model for competing risks, and the loss of DeepHit is a combination of a discrete negative log-likelihood and a ranking loss which is scaled by a constant. Therefore, for DeepHit, three more hyperparameters, namely the combination weight ($\alpha$), the normalizing constant ($\sigma$), and number of discrete time-points, should be further tuned. Following Kvamme et al. [9], we use grid search where the search space is as follow while keeping the network structure unchanged:

| **Hyperparameter** | **Values** |
|---|---|
| $\alpha$ | {0,0.2,0.5,0.8,1} |
| $\sigma$ | {0.1, 0.25, 0.5, 1, 2.5, 5, 10, 100} |
| Num. discrete time-points | {10, 50, 100, 200, 400} |

Table 1: Hyperparameter search space and corresponding values.

For both the simulation and real data studies, we performed a grid search using 100 replicates (either simulated or subsets of real data). For each replicate, we further split the training dataset into a training set and a validation set, explored the predefined hyperparameter space, and calculated the C-index of DeepHit on the validation dataset at each grid point. The hyperparameter combination that achieved the highest average C-index across all 100 replicates was selected as the optimal grid point. This optimal setting was then used for variable selection and model evaluation in subsequent analyses.

**SurvDNN**

To ensure comparability, we used the same hyperparameters stated above, with a bagging size to be 100, and the penalty weight $\lambda$ to be $1 \times 10^{-4}$.

##### Survival Probability Prediction given by Cox-based Models

We can obtain predictions from the relative risk models by estimating the survival function using $\hat{S}(t_j|\boldsymbol{x_{i\cdot}}) = \exp[-\hat{H}(t_j|\boldsymbol{x_{i\cdot}})]$. Following Kvamme et al. [9], for Cox-based models (i.e., Cox, XGboost,DeepSurv, and SurvDNN), we can estimate the cumulative hazards at $t_j$ by:

$$\hat{H}(t_j \mid \boldsymbol{x_{i\cdot}}) = \sum_{T_i \leq t_j} \Delta\hat{H}_0(T_i)\, \exp\big[\hat{g}(T_i, \boldsymbol{x_{i\cdot}})\big]$$

where $\Delta\hat{H}_0(T_i)$ is an increment of the Breslow estimate:

$$\Delta\hat{H}_0(T_i) = \frac{\delta_i}{\sum_{k:T_k \geq T_i} \exp\big[\hat{g}(T_i, \boldsymbol{x_{k\cdot}})\big]}$$

and $\hat{g}(T_i, \boldsymbol{x_{i\cdot}})$ is the (log) relative risk prediction given by Cox-based models.

In order to calculate the integrated Brier score, we calculate $\hat{S}(t_j|\boldsymbol{x_{i\cdot}})$ for each individual $i$ at each unique event time point $t_j$ in the test dataset, and then plug-in all the estimates into the formula of IBS.

[7] Raphael Sonabend and Yohann Foucher. survivalmodels: Models for Survival Analysis, 2022. R package version 0.1.191.

[8] X Mi, B Zou, F Zou, and J Hu. Permutation-based identification of important biomarkers for complex diseases via machine learning models. Nature Communications, 12(1):3008, 2021.

[9] Håvard Kvamme, Ørnulf Borgan, and Ida Scheel. Time-to-event prediction with neural networks and cox regression. J. Mach. Learn. Res., 20:129:1–129:30, 2019.
